## Supplementary Material for "A thorough analysis of the contribution of experimental, derived and sequence-based predicted protein-protein interactions for functional annotation of proteins"

Stavros Makrodimitris [1,2,†], Marcel J.T. Reinders [1,3], Roeland C.H.J. van Ham [1,2]

[1]Delft Bioinformatics Lab, Delft University of Technology, Delft, the Netherlands,
[2]Keygene N.V., Wageningen, the Netherlands and
[3]Leiden Computational Biology Center, Leiden University Medical Center, Leiden, the Netherlands.
[†]To whom correspondence should be addressed.

#### Statistics of PPI networks

The experimental PPI networks of the four species have very different degree distributions, as shown in Figure S1. Table S1 lists the number of total and unique protein-protein interactions (PPIs) that each data source includes for *Saccharomyces cerevisae*, *Escherichia coli*, *Arabidoisis thaliana* and *Solanum lycopersicum.*

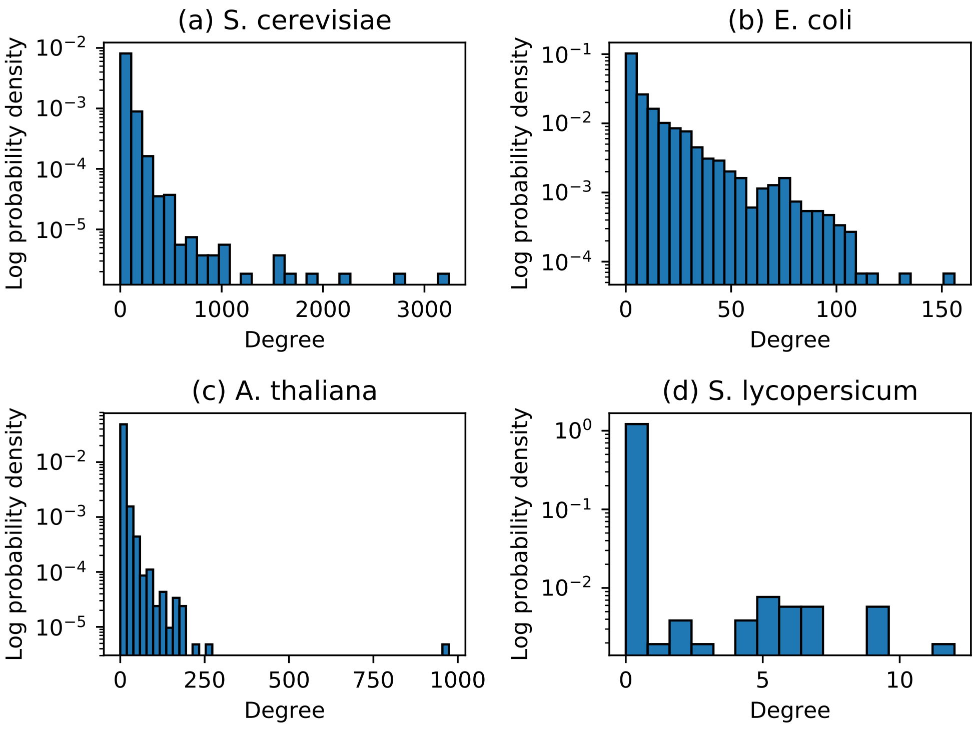


Figure S1: Histogram of node degrees in the experimental PPI networks of yeast (a), *E. coli* (b), arabidopsis (c) and tomato (d). On the $x$-axis node degree and on the $y$-axis the probability density in log scale.

Table S1: Number of interactions added by each data source (rows) in each species (columns). "Database" from STRING was not considered, as it includes protein associations based on known GO annotations

| Data source | *S. cerevisiae* | | *E. coli* | | *A. thaliana* | | *S. lycopersicum* | |
|---|---|---|---|---|---|---|---|---|
| | Total | Unique | Total | Unique | Total | Unique | Total | Unique |
| Physical (BIOGRID) | 97,503 | 51,473 | 9,734 | 5,295 | 21,720 | 10,826 | 57 | 11 |
| Experiments (STRING) | 87,378 | 24,485 | 11,269 | 3,944 | 10,176 | 746 | 0 | 0 |
| Neighborhood (STRING) | 0 | 0 | 3,644 | 638 | 0 | 0 | 0 | 0 |
| Neighborhood transferred (STRING) | 55,410 | 30,241 | 49,167 | 27,128 | 165,627 | 106,821 | 117 | 57 |
| Co-Occurence (STRING) | 2,217 | 498 | 43,114 | 29,663 | 21,283 | 12,968 | 1,014 | 187 |
| Database transferred (STRING) | 0 | 0 | 0 | 0 | 0 | 0 | 0 | 0 |
| Experiments transferred (STRING) | 121,858 | 57,464 | 13,356 | 6,175 | 351,488 | 236,893 | 2,878 | 1,247 |
| Fusion (STRING) | 1,784 | 1,007 | 1,568 | 224 | 2,010 | 671 | 2 | 0 |
| Homology (STRING) | 3,700 | 561 | 1,083 | 414 | 31,983 | 16,734 | 1,858 | 212 |
| Co-Expression (STRING) | 71,525 | 24,635 | 0 | 0 | 285,075 | 217,555 | 0 | 0 |
| Co-Expression transferred (STRING) | 242,228 | 100,849 | 47,897 | 21,878 | 390,727 | 193,802 | 3,029 | 622 |
| Text mining (STRING) | 206,721 | 122,099 | 7,609 | 4,798 | 222,844 | 171,286 | 904 | 595 |
| Text mining transferred (STRING) | 211,409 | 95,945 | 56,745 | 30,854 | 497,495 | 313,994 | 3,712 | 1,412 |

#### Integration of STRING scores

For completeness, we describe the algorithm for integrating STRING scores from $D$ different sources [1]. We use $s_i$ to denote the score (probability) of a particular interaction based on data type $i$. This score is computed by taking into account the prior probability of two proteins interacting ($p$), for which the value $p = 0.041$ is used. When combining multiple data types, this prior score should be only be incorporated once, so the first step is to remove it from all data types (Eq 1):

$$s_{i,noprior} = \frac{s_i - p}{1 - p} \tag{1}$$

Scores from different sources are combined using Eq 2:

$$s_{combined,noprior} = 1 - \prod_{i=1}^{D}(1 - s_{i,noprior}) \tag{2}$$

Finally, the posterior probability is incorporated back into the score by Eq 3, which is the same as Eq 1, but solving for a different variable.

$$s_{combined} = s_{combined,noprior} + p(1 - s_{combined,noprior}) \tag{3}$$

#### *node2vec* parameters and tuning

The *node2vec* algorithm has several parameters that can have great influence on the final node embeddings [2] and therefore the function prediction performance. For each species, we tuned these parameters for the experimental network and for the best combination of experimental and STRING edges separately. To do so, we employed a grid search. Below is a list of the parameters we tuned and the ranges of parameter values we considered:

- The number of random walks per node ([5, 10, 15, 20, 30]),
- The parameter $p$ [2] ([0.1, 0.5, 1.0, 1.5, 3.0]),
- The parameter $q$ [2] ([0.1, 0.5, 1.0, 1.5, 3.0]),
- The length of each random walk ([20, 40, 60, 80, 100, 160]), and
- The dimensionality of the embedding vector ([32, 64, 96, 128, 150, 200, 500]).

For each of the combinations above, we tried the $k$NN and ridge classifiers. The $k$NN also has one tunable parameter, the number of neighbors $k$, for which we considered the values [1, 2, 5, 7, 9, 11, 15, 21, 31, 51]. The parameter $\lambda$ of ridge, which controls the amount of L2 regularization, was tuned among the values [0.0001, 0.001, 0.01, 0.1, 0.5, 1.0, 2.0, 5.0, 10.0].

This amounts to 105,000 classification models per network per species. Each model was evaluated in each of the five cross-validation folds, by further subdividing the training set of each fold into a training (80% of original training set) and a validation set (20% of originial training set). The combination of *node2vec* hyperparameters, classifier and classifier parameter that maximized the validation $F_{max}$ in each fold was subsequently trained on the entire training set (of the corresponding fold) and the resulting trained model was tested on the previously unseen test set.

### Evaluation measures and results

We evaluated using the protein-centric F1 score and Semantic Distance [3]. Both metrics are calculated using hard (0 or 1) predictions and not posterior probabilities. Similar to the CAFA challenges [4], we report the metric value of each metric for the optimal operating point, which justifies the names $F_{max}$ and $S_{min}$. We examined all operating points from 0 to 1 with a step 0.02. The performance of all tested methods is listed in Tables S2 and S3.

Table S2: $F_{max}$ of different function prediction methods (rows) in 4 species (columns) estimated using 5-fold cross-validation. The mean ±the standard deviation is shown.

| | *S. cerevisiae* | *E. coli* | *A. thaliana* | *S. lycopersicum* |
|---|---|---|---|---|
| Naive | 0.31±0.004 | 0.29±0.009 | 0.28±0.005 | 0.23±0.019 |
| BLAST | 0.35±0.004 | 0.43±0.021 | 0.42±0.007 | 0.34±0.025 |
| EXP, GBA | 0.42±0.007 | 0.25±0.011 | 0.19±0.007 | 0.03±0.014 |
| EXP, node2vec | 0.50±0.012 | 0.28±0.011 | 0.23±0.008 | 0.08±0.042 |
| EXP, GBA + BLAST | 0.50±0.007 | 0.45±0.019 | 0.42±0.006 | 0.33±0.025 |
| EXP + STRING, GBA | 0.49±0.005 | 0.46±0.008 | 0.48±0.004 | 0.61±0.045 |
| EXP + STRING, node2vec | 0.59±0.009 | 0.50±0.012 | 0.50±0.005 | 0.61±0.042 |
| EXP + STRING, GBA + BLAST | 0.54±0.006 | 0.58±0.021 | 0.54±0.008 | 0.60±0.047 |
| EXP + SEQ, GBA | 0.33±0.005 | 0.32±0.008 | 0.29±0.004 | 0.33±0.016 |

Table S3: $S_{min}$ of different function prediction methods (rows) in 4 species (columns) estimated using 5-fold cross-validation. The mean ±the standard deviation is shown.

| | *S. cerevisiae* | *E. coli* | *A. thaliana* | *S. lycopersicum* |
|---|---|---|---|---|
| Naive | 34.67±0.75 | 21.41±0.20 | 30.44±0.57 | 17.67±0.77 |
| BLAST | 35.24±0.97 | 16.85±0.85 | 27.66±0.31 | 19.31±0.87 |
| EXP, GBA | 31.17±0.55 | 21.58±0.44 | 30.55±0.61 | 19.39±0.91 |
| EXP, node2vec | 30.05±0.55 | 20.37±0.41 | 28.69±0.59 | 19.37±0.89 |
| EXP, GBA + BLAST | 29.28±0.83 | 15.57±0.79 | 27.75±0.55 | 18.25±0.92 |
| EXP + STRING, GBA | 27.42±0.57 | 17.07±0.22 | 24.12±0.50 | 9.08±0.89 |
| EXP + STRING, node2vec | 25.65±0.48 | 17.23±0.34 | 25.10±0.15 | 9.20±0.77 |
| EXP + STRING, GBA + BLAST | 28.48±0.97 | 12.94±0.60 | 23.00±0.56 | 10.21±1.06 |
| EXP + SEQ, GBA | 34.50±0.74 | 21.02±0.14 | 30.48±0.57 | 17.06±0.71 |

### STRING performance per data source

For every species, we tested all possible combinations of *STRING* data sources and found that certain combinations perform significantly better than others, as certain individual data sources were more informative (Fig S2-S4.). To quantify the contribution of each data source we devised a statistical test described below.

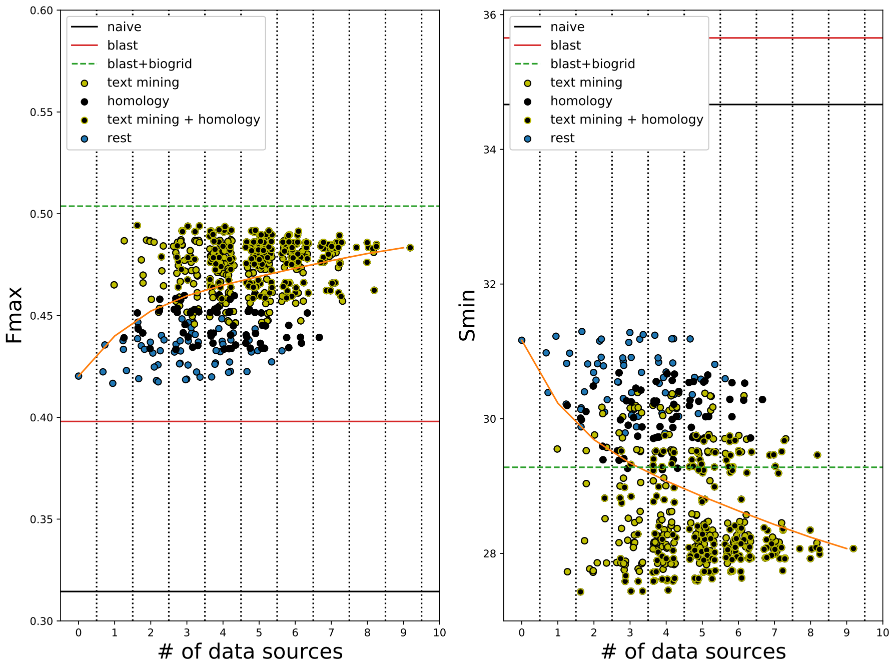


Figure S2: $F_{max}$ (left) and $S_{min}$ (right) in *S. cerevisiae* ($y$-axis) as a function of the number of *STRING* data sources included ($x$-axis). Each dot corresponds to one combination of data sources added to the experimental network. Combinations that include "text mining" and/or "text mining transferred" are shown in yellow, combinations that include "homology" in black and combinations that include both in black with yellow border. The rest of the combinations are shown in blue. To ease visibility, we added a random number in the range [-0.5, 0.5] to each combination of the same number of sources. Zero data sources corresponds to the *EXP* network and the orange line shows the average performance for a specific number of data sources. Horizontal lines denote the performance of the *naive* (black), *BLAST* (red) and the combination of *BLAST* with the *EXP* PPI network (dashed green).

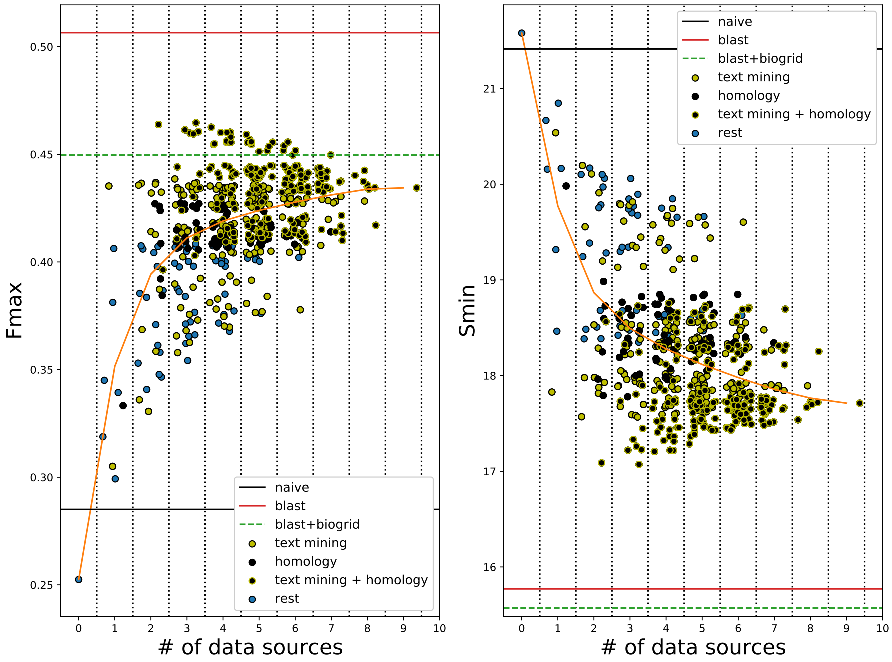


Figure S3: $F_{max}$ (left) and $S_{min}$ (right) in *E. coli* ($y$-axis) as a function of the number of *STRING* data sources included ($x$-axis). Each dot corresponds to one combination of data sources added to the experimental network. Combinations that include "text mining" and/or "text mining transferred" are shown in yellow, combinations that include "homology" in black and combinations that include both in black with yellow border. The rest of the combinations are shown in blue. To ease visibility, we added a random number in the range [-0.5, 0.5] to each combination of the same number of sources. Zero data sources corresponds to the *EXP* network and the orange line shows the average performance for a specific number of data sources. Horizontal lines denote the performance of the *naive* (black), *BLAST* (red) and the combination of *BLAST* with the *EXP* PPI network (dashed green).

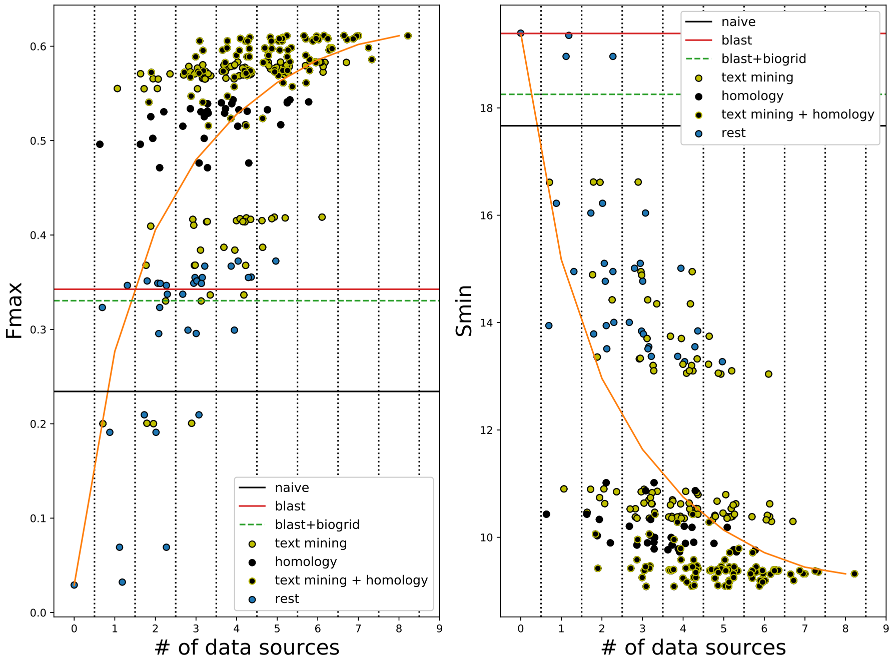


Figure S4: $F_{max}$ (left) and $S_{min}$ (right) in *S. lycopersicum* ($y$-axis) as a function of the number of *STRING* data sources included ($x$-axis). Each dot corresponds to one combination of data sources added to the experimental network. Combinations that include "text mining" and/or "text mining transferred" are shown in yellow, combinations that include "homology" in black and combinations that include both in black with yellow border. The rest of the combinations are shown in blue. To ease visibility, we added a random number in the range [-0.5, 0.5] to each combination of the same number of sources. Zero data sources corresponds to the *EXP* network and the orange line shows the average performance for a specific number of data sources. Horizontal lines denote the performance of the *naive* (black), *BLAST* (red) and the combination of *BLAST* with the *EXP* PPI network (dashed green).

For a species for which *STRING* contains protein associations from $N$ different data sources, there are $2^N - 1$ ways to combine them (in combinations of different lengths). Each data source $i$ is present in $2^{N-1}$ of these combinations (denoted as $P_i$) and absent in the remaining ones (denoted as $A_i$). We define an "irreplaceability" score ($IR(i)$) for $i$ as the increase in the optimal performance of $P_i$ with respect to $A_i$. For $\mathrm{F}_{\max}$, this is formally defined in equation 4

$$IR_F(i) = \max_{c \in P_i} F_{max}(c) - \max_{c \in A_i} F_{max}(c) \quad (4)$$

An $IR_F(i)$ equal to 0, means that the combination that achieves the maximum performance does not include data source $i$, which implies that it made redundant by other sources. A positive value for $IR_F(i)$ means that $i$ is necessary for maximizing *STRING* performance and a negative value that $i$ is creating more errors than correct predictions. Since lower $\mathrm{S}_{\min}$ means better performance, the definition of $IR$ for $\mathrm{S}_{\min}$ is slightly modified, so that a positive $IR$ still denotes that a data source is useful (equation 5).

$$IR_S(i) = \min_{c \in A_i} S_{min}(c) - \min_{c \in P_i} S_{min}(c) \quad (5)$$

We compared the observed $IR_F$ and $IR_S$ values to what would be expected by chance to obtain a measure of statistical significance. To do so, we did a permutation test for which we randomly mixed the elements of $P_i$ and $A_i$ and calculated the $IR_F$ and $IR_S$ values again. We repeated this 10,000 times for each data source to obtain a null distribution of the two statistics and counted the fraction of times the permuted statistic was larger than the observed one. Note that this is a one-sided test, meaning that we only tested whether a data source was significantly irreplaceable, but not whether it was significantly leading to worse performance. We performed 90 tests in total, so we also applied multiple testing correction using the FDR method. The observed test statistics and the corresponding p-values are listed in Tables S4-S5 for yeast, S6-S7 for E. coli, S8-S9 for arabidopsis and S10-S11 for tomato.

Table S4: IRF scores of STRING data sources, corresponding raw p-values and p-values corrected for multiple tests for *S. cerevisiae*

| Data source | $IR_F$ | p-value | corrected p-value |
|---|---|---|---|
| coexpression | -0.0026 | 0.9989 | 1.0 |
| neighborhood transferred | -0.0024 | 0.9933 | 1.0 |
| coexpression transferred | -0.0081 | 1.0 | 1.0 |
| experiments transferred | -0.0078 | 1.0 | 1.0 |
| **textmining** | 0.0154 | $< 10^{-4}$ | $< 0.007$ |
| cooccurence | -0.0004 | 0.8738900 | 1.0 |
| fusion | -0.0002 | 0.75490 | 1.0 |
| textmining transferred | -0.0022 | 0.9715 | 1.0 |
| **homology** | 0.0007 | $< 10^{-4}$ | $< 0.007$ |

Table S5: IRS scores of STRING data sources, corresponding raw p-values and p-values corrected for multiple tests for *S. cerevisiae*

| Data source | $IR_S$ | p-value | corrected p-value |
|---|---|---|---|
| coexpression | -0.1572 | 0.9832 | 1.0 |
| neighborhood transferred | -0.1571 | 0.9707 | 1.0 |
| coexpression transferred | -0.4923 | 1.0 | 1.0 |
| experiments transferred | -0.4663 | 1.0 | 1.0 |
| **textmining** | 1.3206 | $< 10^{-4}$ | $< 0.007$ |
| cooccurence | -0.0105 | 0.7568 | 1.0 |
| fusion | -0.0111 | 0.8798 | 1.0 |
| textmining transferred | -0.1867 | 1.0 | 1.0 |
| **homology** | 0.2914 | $< 10^{-4}$ | $< 0.007$ |

Table S6: IRF scores of STRING data sources, corresponding raw p-values and p-values corrected for multiple tests for *E. coli*

| Data source | $IR_F$ | p-value | corrected p-value |
|---|---|---|---|
| neighborhood | -0.0042 | 0.9695 | 1.0 |
| neighborhood transferred | -0.0198 | 1.0 | 1.0 |
| coexpression transferred | -0.0046 | 0.9962 | 1.0 |
| experiments transferred | -0.0021 | 0.8759 | 1.0 |
| textmining | -0.0045 | 0.9919 | 1.0 |
| cooccurence | -0.0232 | 1.0 | 1.0 |
| fusion | 0.0008 | 0.4949 | 1.0 |
| **textmining transferred** | 0.0303 | $< 10^{-4}$ | $< 0.007$ |
| **homology** | 0.0261 | $< 10^{-4}$ | $< 0.007$ |

Table S7: IRS scores of STRING data sources, corresponding raw p-values and p-values corrected for multiple tests for *E. coli*

| Data source | $IR_S$ | p-value | corrected p-value |
|---|---|---|---|
| neighborhood | -0.1456 | 0.9418 | 1.0 |
| neighborhood transferred | -0.5245 | 1.0 | 1.0 |
| coexpression transferred | -0.2496 | 0.9996 | 1.0 |
| experiments transferred | -0.1348 | 0.8769 | 1.0 |
| textmining | -0.1483 | 0.9839 | 1.0 |
| cooccurence | -0.3763 | 1.0 | 1.0 |
| fusion | 0.016 | 0.4938 | 1.0 |
| **textmining transferred** | 0.6704 | $< 10^{-4}$ | $< 0.007$ |
| **homology** | 0.4951 | $< 10^{-4}$ | $< 0.007$ |

Table S8: IRF scores of STRING data sources, corresponding raw p-values and p-values corrected for multiple tests for *A. thaliana*

| Data source | $IR_F$ | p-value | corrected p-value |
|---|---|---|---|
| coexpression | -0.0297 | 1.0 | 1.0 |
| neighborhood transferred | -0.0225 | 0.96600 | 1.0 |
| coexpression transferred | -0.0324 | 1.0 | 1.0 |
| experiments transferred | -0.0275 | 1.0 | 1.0 |
| **textmining** | 0.0258 | 0.00010 | 0.0052 |
| cooccurence | 0.0001 | 0.50100 | 1.0 |
| fusion | 0.0002 | 0.25070 | 1.0 |
| textmining transferred | -0.0252 | 0.9982 | 1.0 |
| **homology** | 0.0534 | $< 10^{-4}$ | $< 0.007$ |

Table S9: IRS scores of STRING data sources, corresponding raw p-values and p-values corrected for multiple tests for *A. thaliana*

| Data source | $IR_S$ | p-value | corrected p-value |
|---|---|---|---|
| coexpression | -0.9484 | 1.0 | 1.0 |
| neighborhood transferred | -0.7883 | 0.9984 | 1.0 |
| coexpression transferred | -1.2361 | 1.0 | 1.0 |
| experiments transferred | -1.1204 | 1.0 | 1.0 |
| textmining | 0.5247 | 0.0659 | 1.0 |
| cooccurence | -0.0069 | 0.8801 | 1.0 |
| fusion | -0.0066 | 0.7477 | 1.0 |
| textmining transferred | -1.0243 | 1.0 | 1.0 |
| **homology** | 1.9112 | $< 10^{-4}$ | $< 0.007$ |

Table S10: IRF scores of STRING data sources, corresponding raw p-values and p-values corrected for multiple tests for tomato

| Data source | $IR_F$ | p-value | corrected p-value |
|---|---|---|---|
| neighborhood transferred | -0.0002 | 0.98540 | 1.0 |
| coexpression transferred | 0.0019 | 0.00380 | 0.1938 |
| **experiments transferred** | 0.0031 | $< 10^{-4}$ | $< 0.007$ |
| **textmining** | 0.0128 | $< 10^{-4}$ | $< 0.007$ |
| cooccurence | 0.0002 | 0.24900 | 1.0 |
| fusion | 0.0000 | 0.75380 | 1.0 |
| **textmining transferred** | 0.0254 | $< 10^{-4}$ | $< 0.007$ |
| **homology** | 0.0281 | $< 10^{-4}$ | $< 0.007$ |

Table S11: IRS scores of STRING data sources, corresponding raw p-values and p-values corrected for multiple tests for tomato

| Data source | $IR_S$ | p-value | corrected p-value |
|---|---|---|---|
| neighborhood transferred | -0.0029 | 0.93930 | 1.0 |
| coexpression transferred | -0.0812 | 1.0 | 1.0 |
| experiments transferred | -0.2280 | 1.0 | 1.0 |
| **textmining** | 0.1536 | $< 10^{-4}$ | $< 0.007$ |
| cooccurence | -0.0085 | 0.9875 | 1.0 |
| fusion | 0.0000 | 0.75470 | 1.0 |
| **textmining transferred** | 0.2691 | $< 10^{-4}$ | $< 0.007$ |
| **homology** | 1.2114 | $< 10^{-4}$ | $< 0.007$ |

#### Removing most informative data sources

Table S12: Changes in $F_{max}$ of the combined *EXP+STRING* network, when removing text mining and or homology edges

| Data sources | *S. cerevisiae* | *E. coli* | *A. thaliana* | *S. lycopersicum* |
|---|---|---|---|---|
| All | 0.49±0.005 | 0.46±0.008 | 0.48±0.004 | 0.61±0.045 |
| No text mining/text mining transferred | 0.46±0.006 | 0.43±0.013 | 0.45±0.007 | 0.54±0.053 |
| No homology | 0.49±0.004 | 0.44±0.011 | 0.43±0.006 | 0.58±0.044 |
| No text mining/text mining transferred/homology | 0.45±0.006 | 0.41±0.010 | 0.35±0.004 | 0.37±0.042 |

Table S13: Changes in $S_{min}$ of the combined *EXP+STRING* network, when removing text mining and or homology edges

| Data sources | *S. cerevisiae* | *E. coli* | *A. thaliana* | *S. lycopersicum* |
|---|---|---|---|---|
| All | 27.42±0.57 | 17.07±0.22 | 24.12±0.50 | 9.08±0.89 |
| No text mining/text mining transferred | 29.24±0.64 | 17.78±0.43 | 24.65±0.59 | 9.72±1.16 |
| No homology | 27.72±0.57 | 17.57±0.26 | 26.03±0.51 | 10.30±0.84 |
| No text mining/text mining transferred/homology | 29.79±0.63 | 18.35±0.22 | 28.47±0.47 | 13.27±0.65 |

### PIPR training and results

Table S14: Effect of hyperparameters in training of PIPR for predicting yeast PPIs from the BIOGRID database.

| **Optimizer** | **Learning Rate** | **Validation loss** | **Validation accuracy** |
|---|---|---|---|
| ADAM | 0.001 | 0.507 | 0.771 |
| ADAM | 0.0001 | 0.540 | 0.754 |
| SGD + learning rate scheduler | 0.01 | 0.525 | 0.751 |
| SGD + learning rate scheduler | 0.001 | 0.547 | 0.737 |

The originally trained PIPR model in yeast generalized poorly in Arabidopsis (accuracy of 51% on a balanced dataset). Therefore, we chose to train PIPR on Arabidopsis by using the original trained model as initial conditions. We trained using Stochastic Gradient Descent with learning rate 0.001 and early stopping based on the validation loss with patience of 40 epochs. As validation set, we randomly selected 10% of the data and as loss function the binary cross-entropy. We did not use RMSprop optimizer, which was used by the authors, as it produced unstable results.

Table S15: Comparison of function prediction performance in arabidopsis and tomato of edges predicted with PIPR trained in either yeast or arabidopsis

| Train species | Test species | $F_{max}$ | $S_{min}$ |
|---|---|---|---|
| *S. cerevisiae* | *A.thaliana* | $0.29 \pm 0.004$ | $30.48 \pm 0.57$ |
| *A. thaliana* | *A.thaliana* | $0.29 \pm 0.006$ | $30.16 \pm 0.29$ |
| *S. cerevisiae* | *S. lycopersicum* | $0.33 \pm 0.016$ | $17.05 \pm 0.70$ |
| *A. thaliana* | *S.lycopersicum* | $0.35 \pm 0.017$ | $16.24 \pm 0.66$ |
